# MCPdb: The Bacterial Microcompartment Database

**DOI:** 10.1101/2021.01.09.426059

**Authors:** Jessica M. Ochoa, Kaylie Bair, Thomas Holton, Thomas A. Bobik, Todd O. Yeates

**Author notes:** Corresponding author; Contact information: Todd O. Yeates, UCLA Department of Chemistry and Biochemistry Los Angeles, CA 90095.

## Abstract

Bacterial microcompartments are organelle-like structures composed entirely of proteins. They have evolved to carry out several distinct and specialized metabolic functions in a wide variety of bacteria. Their outer shell is constructed from thousands of tessellating protein subunits, encapsulating enzymes that carry out the internal metabolic reactions. The shell proteins are varied, with single, tandem and permuted versions of the PF00936 protein family domain comprising the primary structural component of their polyhedral architecture, which is reminiscent of a viral capsid. While considerable amounts of structural and biophysical data have been generated in the last 15 years, current resources present challenges for understanding the functional and structural properties of microcompartments (MCPs) and their diversity. In order to make the remarkable structural features of bacterial microcompartments accessible to a broad community of scientists and non-specialists, we developed MCPdb: The Bacterial Microcompartment Database (https://mcpdb.mbi.ucla.edu/). MCPdb is a comprehensive resource that categorizes and organizes known microcompartment protein structures and their larger assemblies. To emphazise the critical roles symmetric assembly and architecture play in microcompartment function, each structure in the MCPdb is validated and annotated with respect to: (1) its predicted natural assembly state (2) tertiary structure and topology and (3) the metabolic compartment type from which it derives. The current database includes 134 structures and is available to the public with the anticipation that it will serve as a growing resource for scientists interested in understanding protein-based metabolic organelles in bacteria.

## INTRODUCTION

Bacterial microcompartments (MCPs or alternatively BMCs), are supramolecular structures found in approximately 20% of bacteria across numerous phyla [1,2]. These giant protein-based structures have evolved to serve organelle-like functions, with different MCP types encapsulating distinct enzymes in order to carry out specialized metabolic processes in a sequestered environment within the cell interior [3–6]. MCPs are known to carry out diverse metabolic processes; their unifying feature is that they provide a mechanism for bacteria to perform certain multistep reactions in a way that retains metabolic intermediates inside the MCP. The co-localization of sequentially acting enzymes housed inside the MCP helps optimize metabolic flux while limiting alternative side reactions. Importantly, MCPs help prevent the efflux of toxic and/or volatile intermediates into the cytosol [3,7,8]. Bacterial microcompartments can be broadly classified into two major categories: carboxysomes and metabolosomes. Carboxysomes are the founding members of the MCPs, and the simplest representatives. They enhance CO_2_ fixation in bacteria by encapsulating two sequentially acting enzymes -- carbonic anhydrase and ribulose-1,5-bisphosphate carboxylase/oxygenase (RuBisCO) [4,9,10]. Bicarbonate (in addition to ribulose-bisphosphate) is the substrate entering the carboxysome via diffusion across the shell; CO_2_ is the key intermediate, which is produced by carbonic anhydrase and must be consumed by RuBisCO prior to escape. By contrast, metabolosomes use an assortment of key enzymes to metabolize a variety of substrates including 1,2-propanediol for the propanediol utilization (PDU) MCP, and ethanolamine for the ethanolamine utilization (EUT) [4,7,8,10]. Other microcompartments utilize glycyl-radical chemistry (GRM MCPs) and can be further divided into subclasses based on their substrates and signature enzymes, including the glycyl-radical propanediol (Grp) MCP, the choline utilization (Cut) MCP and an additional GRM type that utilizes fucose and rhamnose [11–15].Lastly, there are MCPs that have been more recently discovered whose metabolic functions are still emerging, including the RMM/Aaum MCP and the Etu MCP. Several recent structures of both BMC and BMV proteins have been determined for an MCP first called RMM (for *<u>R</u>hodoccus and <u>M</u>yonacterium <u>M</u>icrocompartment)* and then renamed Aaum (for its apparent role in amino acetone utilization [1,5,14,16,17]. Additionally, the Etu MCP, or the ethanol utilization microcompartment, has been observed in *Clostridium kluyveri* and has had one of its shell proteins characterized [18,19].

Despite their functional diversity, bacterial microcompartments are now understood to be structurally similar. Constructed entirely of proteins, the outer microcompartment shell is composed of thousands of homologous tessellating shell proteins belonging to the BMC protein family [20–22], whose structures were first elucidated in 2005 [23,24]. The canonical BMC protein domain (Figure 1) oligomerizes to form hexameric disks with central pores for the (presumably) diffusive influx of metabolic substrates and the efflux of products. The hexameric disks pack laterally to form the nearly flat facets of the intact shell, while pentameric BMV proteins form the vertices of these large, polyhedral structures (Figure 1) [15,20]. Any single microcompartment type is composed of multiple paralogs of the BMC protein, with different paralogs offering distinct structural properties. This roughly 100-amino acid domain (Pfam PF00936) remains the primary key for exploring and discovering new types of microcompartments, and has been extensively studied and characterized by numerous groups [15,21,22,25–32]. Structural studies have revealed major topologically distinct variations of the BMC protein domain. The most abundant, canonical form is the BMC-H shell protein, which contains a single BMC domain and forms a cyclic homohexamer (Figure 2A) [23]. An alternate topological form of lesser-understood function occurs in the form of permuted BMC proteins [29]. These contain a single, essentially intact BMC domain that possesses a circular permutation. This circular permutation results in a reordering of the amino acid sequence but a similar overall BMC protein fold (Figure 2B), with some of these structures revealing a high degree of flexibility and symmetry-breaking [28,31]. The BMC-T (*T* stands for tandem) category of proteins consists of two tandem repeats of the BMC domain. BMC-Ts are cyclic trimers that form pseudohexamers (Figure 2C) whose overall shape closely resemble a canonical BMC hexamer [28,33,34]. Further variations exist within the BMC-T type, according to whether the individual domains exhibit circular permutation. In some cases, BMC-T shell proteins have been shown to undergo large conformational changes between closed and open pore states, with critical implications for regulated transport [28,33–36]. Some BMC-Ts and even some BMC-H shell proteins have been found to bind iron-sulfur clusters in their central pores [12,30,37]. Finally, BMV proteins (sometimes referred to as BMC-P) are cyclic homopentamers that form the vertices of bacterial microcompartments (Figure 2D) [15,20,38,39]. These are based on the Pfam03319 protein domain, which is entirely unrelated in sequence, structure and evolution from the BMC protein domain family. The sophisticated mechanistic features of MCPs emphasize their qualification as true organelles in bacteria, built from proteins rather than a lipid bilayer.

**Figure 1.**
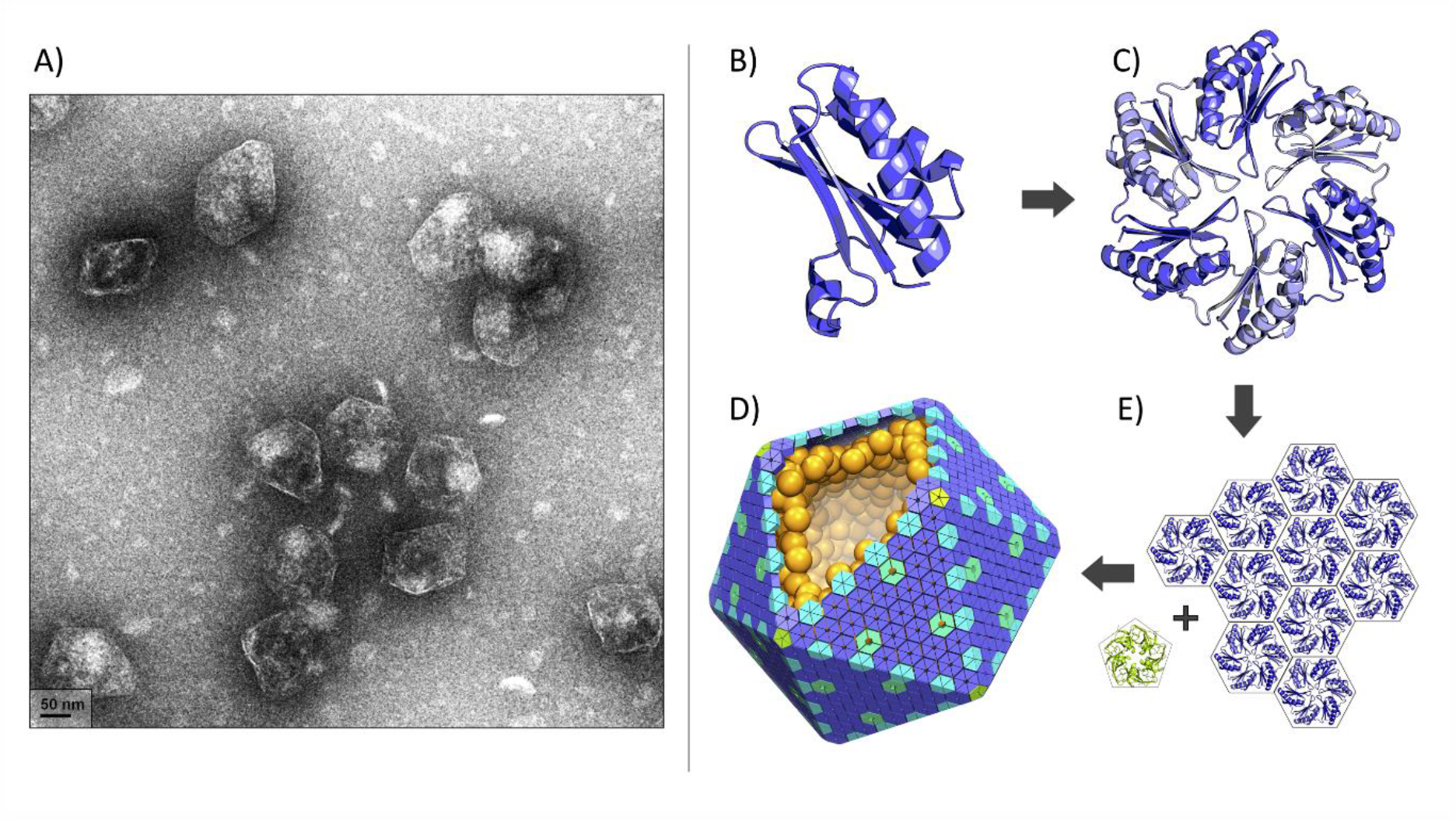
Bacterial microcompartments (MCPs) are large proteinaceous assemblies that function as metabolic organelles. Figure adapted from Ochoa 2020 [31]. (A) Negative stain electron micrograph of purified Pdu MCPs (scale bar: 50 nm). (A) MPC shells are assembled primarily from proteins belonging to the BMC family (A), which are hexameric or trimeric pseudohexamers (B). (C) Hexameric and pseudohexameric BMC shell proteins pack laterally to form the facets while pentameric BMV proteins (yellow green) of unrelated structure form vertices (C). (D) An idealized model of a microcompartment with external shell proteins and encapsulated enzymes. Most natural MCP shells are not as geometrically regular as depicted here by the icosahedral architecture.

**Figure 2.**
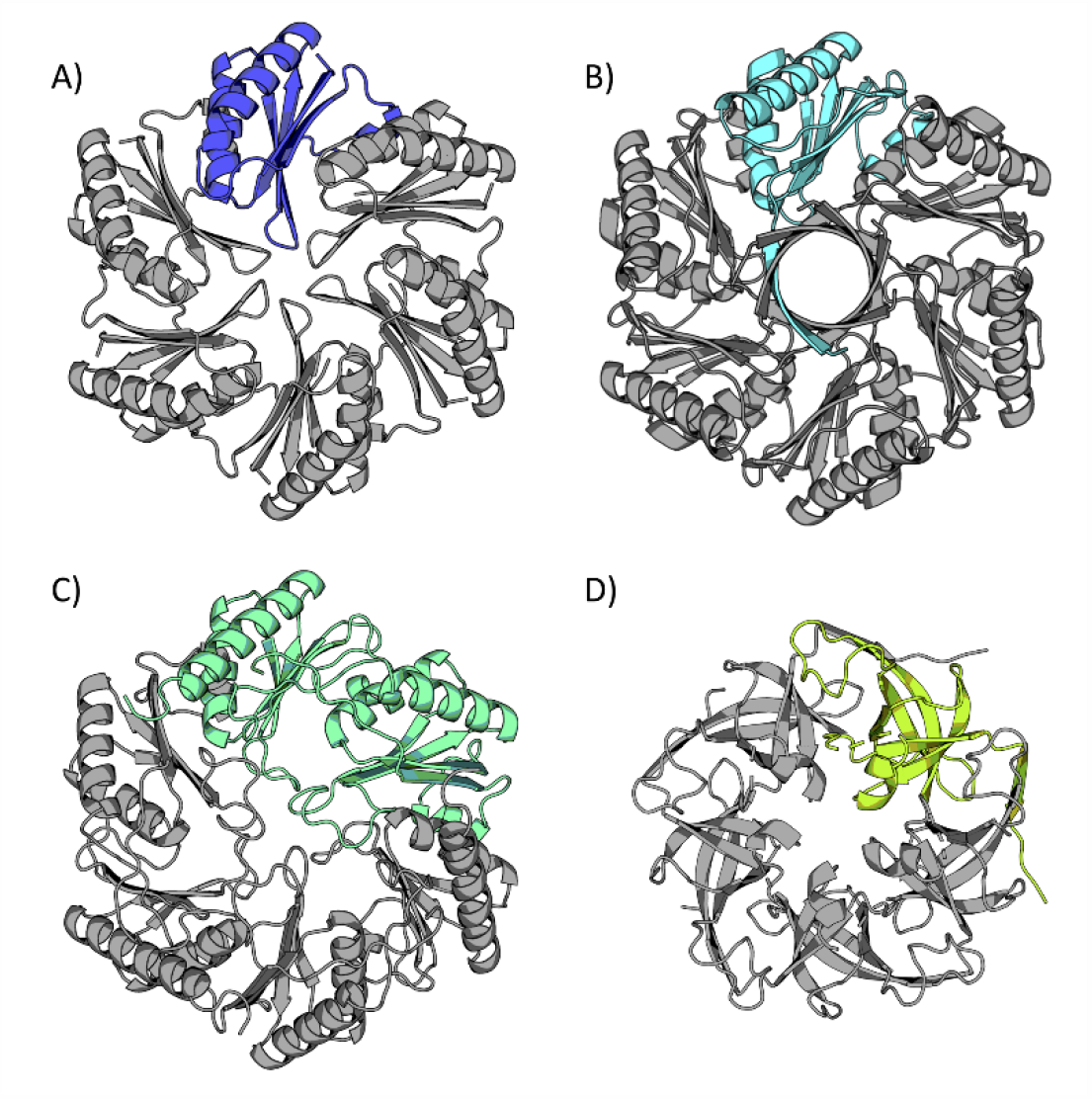
Cartoon representations of four bacterial microcompartment shell proteins. A single monomer is highlighted and presented in the context of the biological assembly. (A) A representative hexameric BMC shell protein (BMC-H) (PDB 2EWH) [24]. (B) A representative permuted BMC shell protein (PDB 6XPI) [31]. (C) A representative trimeric BMC shell protein (BMC-T) (PDB 3I82) [28]. (D) A representative BMV shell protein (PDB 4I7A) [15].

Notwithstanding their wide distribution and the extensive investigation into their outer shells, microcompartments remain only partially understood. To date, more than 120 bacterial microcompartment-related structures have been characterized and deposited in the Protein Data Bank (PDB) (Figure 3). Various items of information about each structure – organism, amino acid sequence, functional name, etc. – are generally available, but other critical insights about structure and function are difficult to sort out from the raw data as it is typically presented, and this challenge is especially true for non-experts that have minimal familiarity with the PDB protein structure database. Because understanding quaternary structure – i.e. protein assembly states – is especially critical to understanding elements of MCP function, we viewed the challenges associated with identifying natural assembly forms as a major barrier for novices trying to generate and understand the natural biological forms of MCP shell proteins. We have also addressed MCP-specific aspects of form and function that are not easily discerned from raw structure files. At present, articulating metabolic MCP type requires some knowledge, and this is provided by curation. Discrimination of diverse topological forms of the BMC proteins is also provided. This is usually non-obvious from raw structural data files, and these structural variations often relate to important properties of the pores (e.g. ‘open’ or ‘closed’), which are routes for metabolite transport.

**Figure 3.**
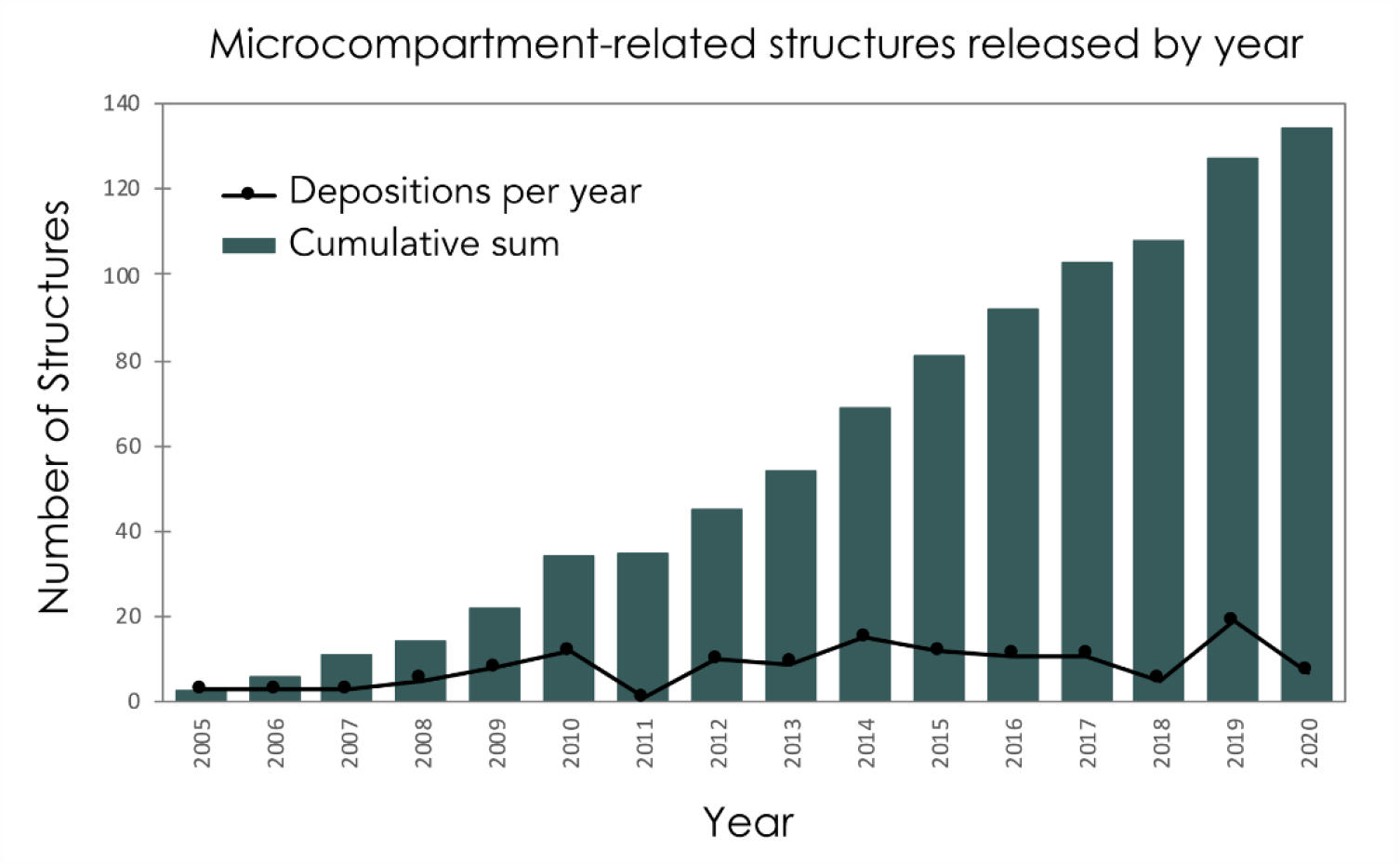
Growth over time of known microcompartment-related structures. There are currently 134 microcompartment and encapsulin-related protein structures deposited in the PDB.

A growing appreciation of the uniqueness and biological importance of MCPs, an expanding body of data on their shell proteins, and current paucity of systematic annotation motivated the development of a centralized database to address these knowledge gaps. Making bacterial microcompartments more accessible to not only structural experts but to a broader scientific audience should help advance this growing field of biology. Here, we describe the development of a novel database, MCPdb: The Bacterial Microcompartment Database (https://mcpdb.mbi.ucla.edu/). While metabolic compartments based on the common BMC protein architecture are the main focus of this database, we also make connections to somewhat related areas of bacterial cell biology by including basic information on encapsulins, a somewhat lesser-studied family of nanocompartments. We collected all known bacterial microcompartment protein structures and assembled a novel online tool that provides users with simplified searching capabilities, structural and biophysical annotations, validated structures, and multiple visualization avenues for examining microcompartment biological assemblies.

## MATERIALS and METHODS

### Data Collection and Curation

MCPdb is built by extracting relevant data from the Protein Data Bank [40] and UniProt [41]. We compiled a list of 134 bacterial microcompartment and encapsulin-related structures. A preliminary list of relevant structures was obtained using keyword searches through the PDB web server (https://www.rcsb.org/). An initial search of the term *microcompartment* yielded 112 structures that required manual validation and verification, resulting in a total of 98 structures related to MCPs. In order to curate a more comprehensive list, we performed searches of structures using the amino acid sequences of representative BMC-H, BMC-T, BMV and permuted BMC structures. With the addition of several structures from (the unrelated) encapsulins, MCPdb presently consists of 134 structures (Figure 4). We performed an HTML-based query to collect relevant information including structure resolution, deposition authors and citations (Figure 5). After generating a master list of PDB IDs, we curated relevant amino acid data from UniProt. A total of 75 unique UniProt IDs gives rise to the 134 separate PDB structures.

**Figure 4.**
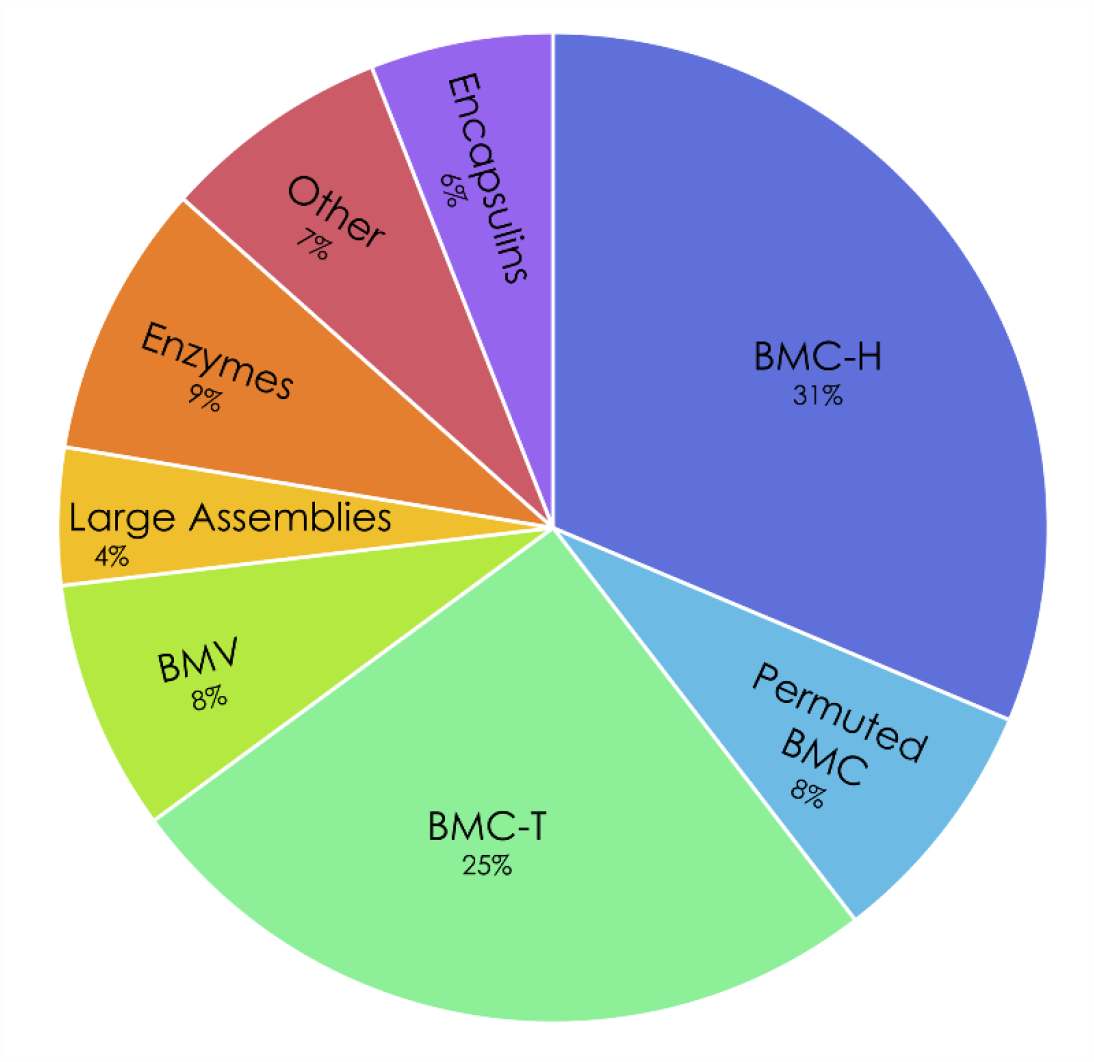
Distribution of protein structure types in the MCPdb. More than 70% of all structures are microcompartment BMC shell protein hexamers or trimers (BMC-H, permuted BMC, BMC-T) or pentamers (BMV), with larger icosahedral assemblies comprising 4%, internal enzymes comprising 9% and other microcompartment associated proteins comprising 7%. Encapsulin structures make up the remaining 6% of the MCPdb.

**Figure 5.**
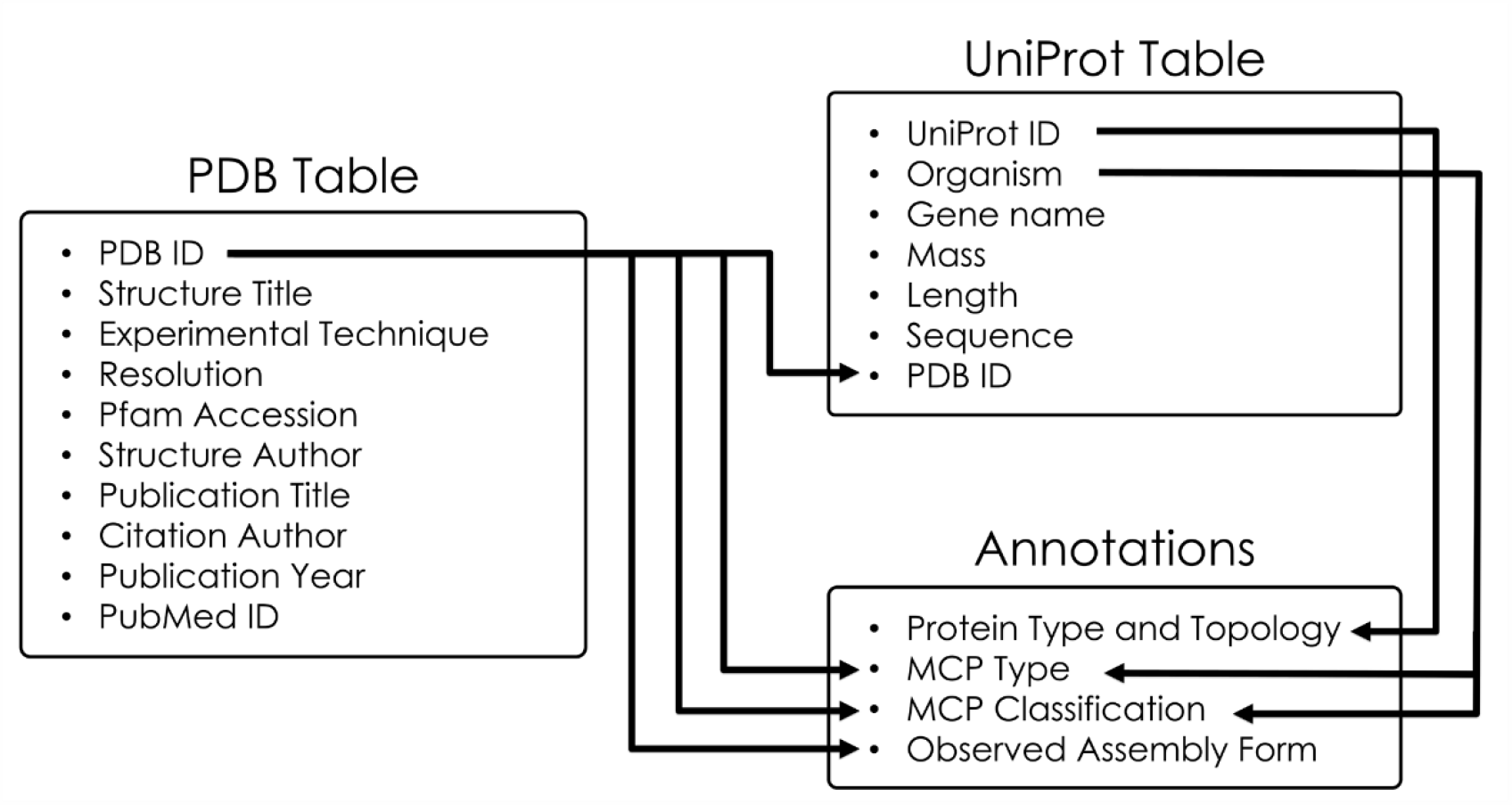
Data sources and annotations for entries in the MCPdb. Key structural information from the PDB as well as associated protein information from UniProt are used to describe each entry. The PDB is the primary link to UniProt IDs. The PDB data file aprovides information about the observed assembly form for the protein, and UniProt provides information from which the protein topology (e.g. circular permutations and domain duplications) can be discerned, Those data sources and the iterature are used to annotate the MCP functional type and subclassification.

Upon collecting all relevant data from the PDB and UniProt, we assigned a series of classifications and annotations to each structure. While individual PDB IDs were used as a key for pertinent structural information, the UniProt IDs were used to provide additional protein details (Figure 5). Each structure in the database has been assigned an *MCP Type, MCP Classification, Protein Type and Topology*, and *Observed Assembly Form* (Figure 6). *MCP Type* broadly categorizes each structure as a carboxysome, a metabolosome or an encapsulin, while *MCP Classification* provides more details about the microcompartment based on its metabolic function, distinguishing between alpha/beta carboxysomes, and the different metabolosome types including the propanediol utilization MCP, ethanolamine utilization MCP and others. We likewise categorized each structure by intrinsic characteristics including *Protein Type and Topology* and *Observed Assembly Form*. While the *Protein Type and Topology* are inherent and describe the type of protein for a given structure (*i*.*e*. BMC-H, BMC-T, BMV, etc.), the *Observed Assembly Form* describes the experimental crystal packing (in some case) and presumptive quaternary architectures.

**Figure 6.**
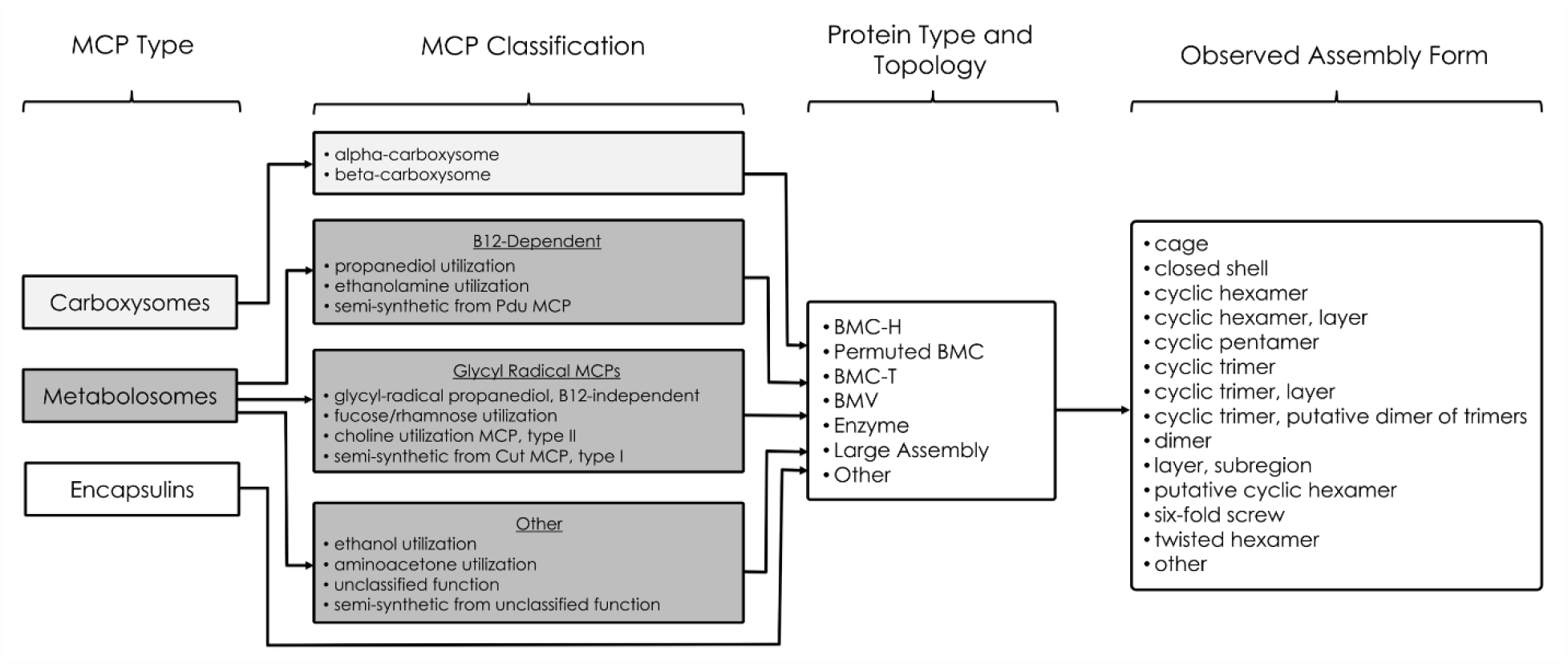
MCPdb entry annotations. MCP Type indicates the broad metabolic category. MCP Classification further distinguishes between the different metabolic subtypes of carboxysomes and metabolosomes. Protein Type and Topology describe properties inherent to the protein tertiary structure. Lastly, Observed Assembly Form describes protein quaternary characteristics of the experimentally described structure.

SQL tables were created to link PDB IDs, UniProt IDs and annotations. In order to construct our database, we utilized a Linux server running Ubuntu 20.04 LTS and MySQL version 5.7. CSV files of the PDB data, UniProt data and annotations were converted into SQL tables with the construction of a linker table to join the tables in the query and to establish the one-to-many relationships between PDB IDs and UniProt IDs (Figure 7). One UniProt can be associated with numerous PDBs (i.e. if the same protein has been structurally characterized in the context of multiple experiments) and one PDB can be associated with numerous UniProts (i.e. if the structure characterized is comprised of proteins of more than one identity). We then generated a series of PHP scripts to query the data and populate our website content. Structures on MCPdb are organized and called by their four-character PDB ID.

**Figure 7.**
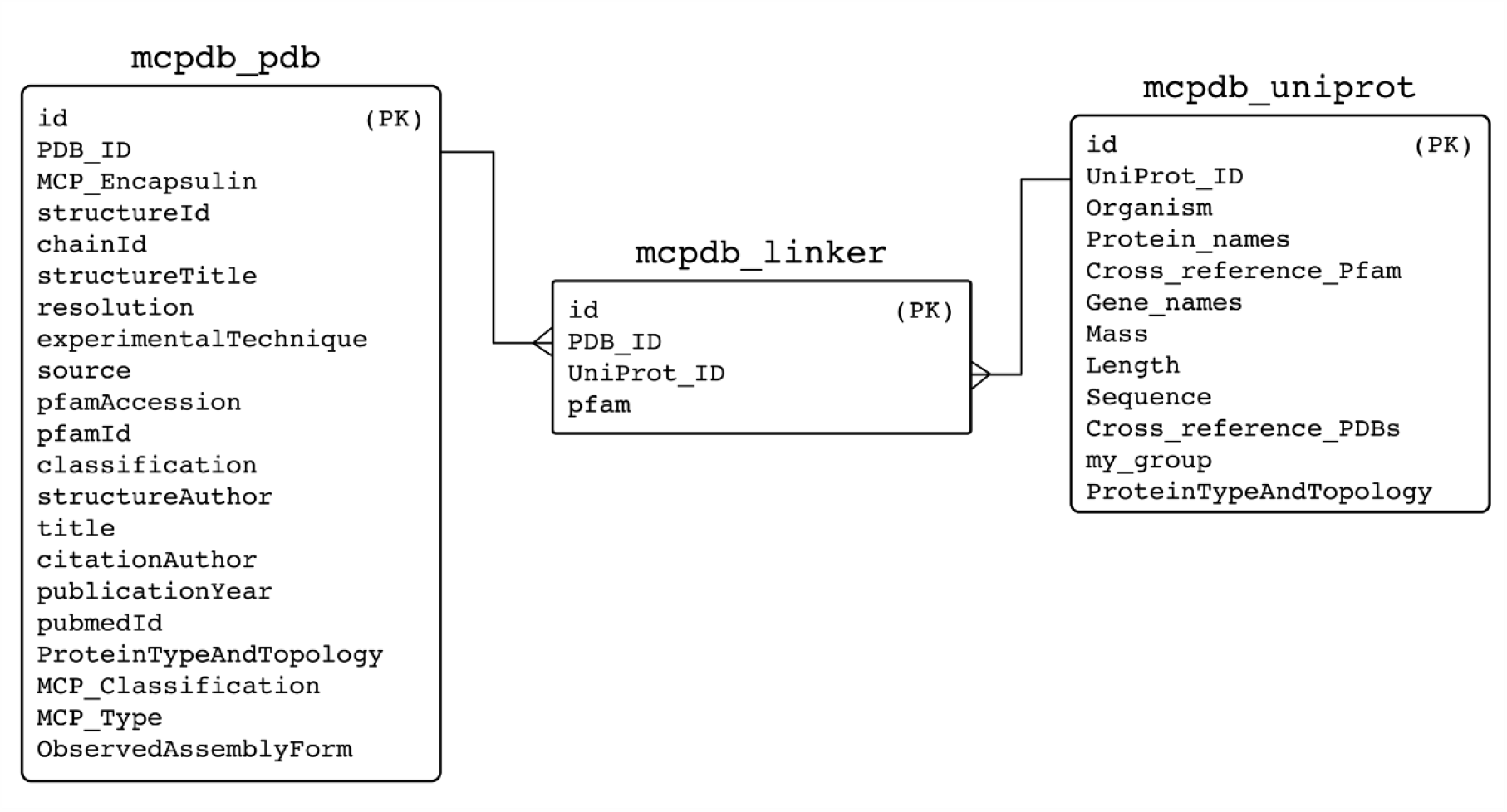
Entity relationship diagram of the MCPdb as a MySQL database. Boxes show the primary data sources and the linker table used to join the tables in the queries. Primary keys (PK) have been identified.

### File Curation and Preparation

In order to construct a centralized microcompartment database, we extracted and compiled relevant files including PDBs, biological assemblies, and FASTA amino acid sequence files with the goal of providing these files to the end user. We also sought to provide users with numerous modes of interacting with each structure. To appeal to experts and novices alike, we incorporated: (1) an interactive 3D viewer for rapid structure interrogation, (2) ready-to-use PyMol graphics sessions for streamlined figure preparation, and (3) images for quickly viewing and interpreting structures while browsing the database.

All files are housed on our permanent institutional web server using the PDB ID as the primary identifier. With the master list of 134 PDB IDs, we utilized a wget command to pull atomic coordinates of all structures in the form of .pdb and .cif files onto our Linux server, which are available to our users as downloads. We were also able to retrieve nearly 60% of the correctly named and trimmed biological assemblies using the program PISA [42]. The biological assemblies that were generated by PISA and migrated to our server were validated for accuracy. The remaining structures whose biological assemblies could not be successfully generated with PISA required manual intervention; the need for this step highlights one of the key utilities of the database. We used PyMol to rename chains in a logical manner and generate new .pdb files. These validated biological assemblies have been cleaned to exclude most small molecules judged to not reflect biological function (e.g. crystallization buffer molecules, etc.). In a few select cases, the natural biological assembly form of a BMC protein remains uncertain (some BMC-T trimers tend to occur in structural studies in the form of two stacked disks). In those cases, users can access alternate assembly forms. MCPdb also provides relevant sequence information in the form of .fasta and .txt files. FASTA-formatted sequences for each structure were retrieved from the PDB; these reflect the actual sequence of the experimentally characterized protein, which can include mutations and the addition of protein purification tags. The native, unmodified protein amino acid sequences (.txt) are extracted from the UniProt data using a PHP query. These files are also available as downloads (Figure 8).

**Figure 8.**
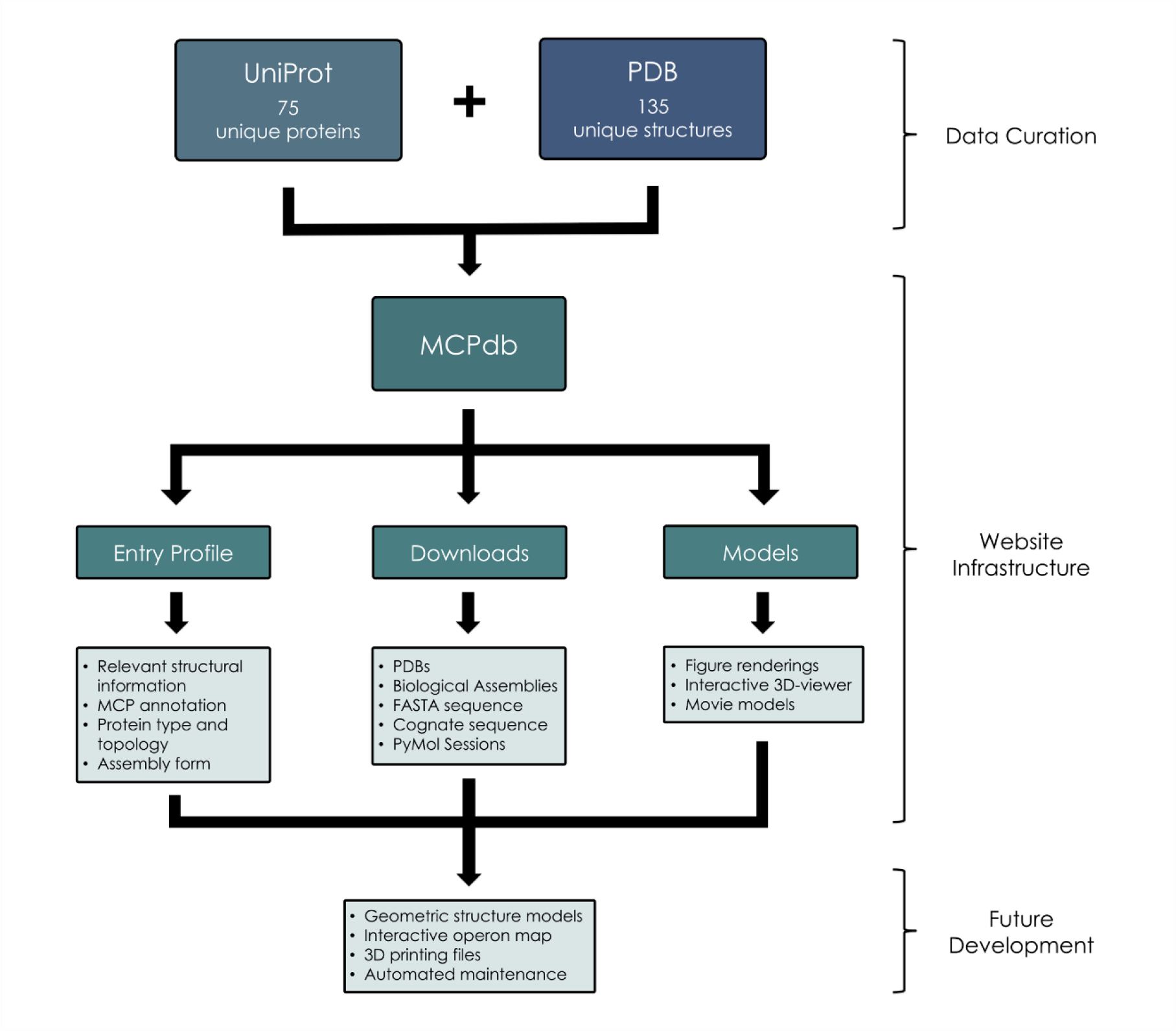
Flow chart depicting data curation and generation, website infrastructure and goals for future development.

We incorporated an interactive 3D viewer that enables users to dynamically engage with most of the MCPdb structures without the use of additional molecular visualization software (Figure 8). The mutation position imaging toolbox (MuPIT) is a browser-based visualization application originally designed for novice structure investigators [43]. By integrating this unmodified software into our database, we provide users the opportunity to quickly visualize structures of interests on desktop and mobile-based browsers. Users may view ribbon, line and stick models of each structure. About 10% of structures were too large for the interactive viewer (some contain as many as 540 protein chains), in which cases the server offers movies (created in PyMol) that dynamically change views and toggle through ribbon and surface renderings of the structure of interest. The short movies of these structures are played in the browser and can also be downloaded and saved locally.

Additionally, we provide ready-to-use PyMol session files (.pse) as optional downloads. These are functional even for the largest of the structures. After curating and manually validating our library of biological assemblies, we prepared a series of PyMol sessions (Figure 8). For each structure, we provide users with a cartoon and surface representation of each structure. Structures are colored such that users can rapidly distinguish between multiple polypeptide chains. Surface representations are semi-transparent for easy viewing. We have also pre-loaded short movies so that users are immediately presented with a rotating view of the selected structure upon launching the PyMol session. Rendering surface representations of large structures, including cages and closed shells, is computationally taxing and can crash PyMol under some computer user configurations. To overcome these challenges, we employed various lesser-known PyMol strategies. By reducing the surface quality and altering the Gaussian resolution option in the Fourier filtering representation prior to generating isosurface maps, we were able to create surface representations, even for the largest structures, which are visually informative while requiring significantly reduced computing power. Our uniform PyMol sessions create an effortless way for novice PyMol users to interact with each of the 134 structures and a simple way of preparing accurate and illustrative figures. Lastly, we generated a series of three figure-ready images (.pngs) for users to scroll through as they are browsing a structure on MCPdb (Figure 8). Based on specifically crafted PyMol sessions, we exported a series of views as .pngs and added these as sliders to each entry in the MCPdb. We additionally created and included N to C-terminus rainbow-colored cartoon diagrams of the asymmetric unit of each structure in the image slider.

### Website Construction

Following database curation, we generated a user-friendly browser interface. The MCPdb infrastructure was created using WordPress, HTML, CSS and JavaScript. We utilized the WordPress graphical user interface (GUI) to build the landing page and accessory information pages. We used HTML, CSS and JavaScript to generate a template page that displays select information for each structure. We replicated and auto-populated data fields in this template for each of the 134 structures using a PHP script. We also created a series of queries to provide our users with seamless and intuitive search features. Infrastructure for simple searches based on key words and more complex filtering searches were also created using PHP.

## RESULTS and DISCUSSION

### Database Description

MCPdb is available at https://mcpdb.mbi.ucla.edu, a permanent institutional URL managed by the UCLA-DOE Institute for Genomics and Proteomics. MCPdb was created in order to compile and consolidate structures related to MCPs, and the several encapsulin structures that are known at this time. More importantly, MCPdb was designed to provide users with readily available structural and biophysical annotations as well as validated biological assemblies. The current version of our database pulls together 134 structures from the PDB (comprising 75 unique UniProts) and a collection of curated files and utilities that are available for in-browser viewing and download (Figure 8). Downloads include PDB files, biological assembly structures (pdb format), Fasta files for biological and experimental protein sequences, and ready-to-use PyMol session files. Users can view rendered images of each structure or interact with them in 3D within the browser. In alignment with our philosophy of introducing new users to the field, MCPdb is freely available and optimized for accessibility on desktops, tablets and mobile devices (Figure 9).

**Figure 9.**
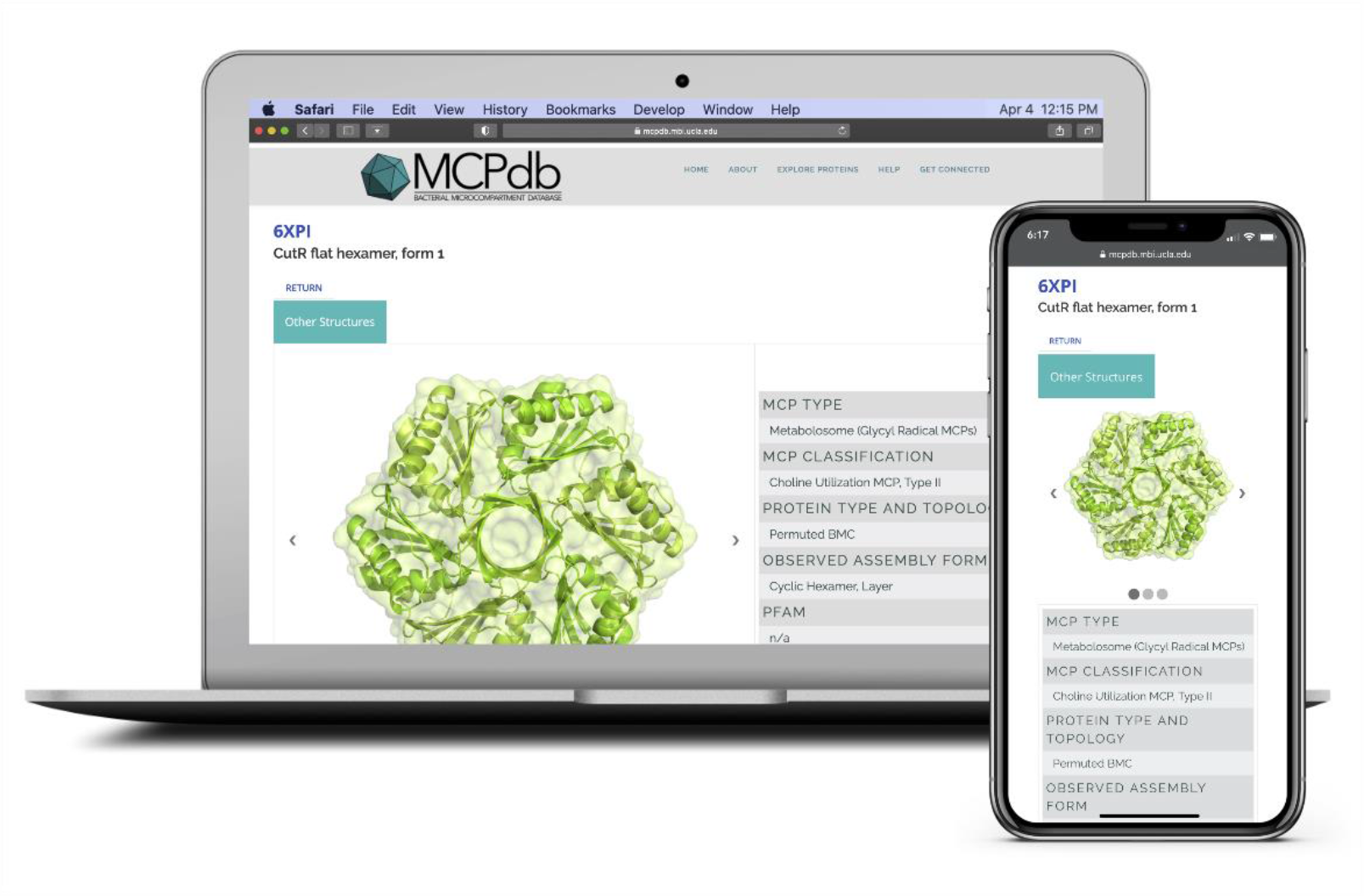
Screenshot and example of a structure profile on the MCPdb interface. MCPdb has been optimized for use on desktops, tablets and mobile devices.

### Web interface

MCPdb provides a simple and interactive framework for users to explore bacterial microcompartments and encapsulins. Upon navigating to the home page, users are presented with a brief database description and provided with links that navigate to a summary page, search page and a quick-start guide. As users explore MCPdb, they are introduced to high-level information about MCPs and their characteristic shell proteins. As they navigate to an individual entry page, users are provided with images of the structure and relevant annotations including *MCP Type, MCP Classification, Protein Type and Topology* and *Observed Assembly Form* (Figure 9). Users can scroll down for additional information related to the structure and authorship, download validated structure files and ready-made PyMol sessions or view the structure in 3D within their browser. We additionally provide a *Get Connected* page to allow users to request assistance and provide feedback.

### Comparison to Other Databases

The MCPdb combines data available from other sources, including the PDB [30] and UniProt [31], with curation and substantial post-processing. The various curation and postprocessing protocols add considerable value compared to currently available data repositories. Presentation of correct biological assembly states is often a challenge for structures obtained by crystallographic methods, and as noted above this is a critical aspect of interrogating MCP structure and function. Vital information, and search capacity, is also provided concerning metabolic function types and unique topological features in the BMC protein family; these features relate to functional differences in their assembly and their roles in molecular transport. There are some parallels between MCPs and viral capsids, and indeed the need for a database for curating biological assembly forms for viral capsids was recognized some years ago with the development of the VIPERdb database [44]. The MCPdb answers the corresponding need for bacterial microcompartments.

Curation has been applied to remove complicating accessory data (e.g. bound buffer molecules, conflicting polypeptide chain names, etc), which might otherwise confuse non-expert users. The integration with multiple modes of visualization, tailored where necessary according to size, will facilitate the graphical display and dissemination of information on these special biological systems. Attention has been given to providing simple methods of display to serve the broadest community of users.

## CONCLUSION and FUTURE PROSPECTS

The MCPdb currently houses 134 microcompartment protein and encapsulin-related structures. Access to validated biological structures as well as structural and biophysical annotations is necessary for well-informed scientific investigation surrounding MCPs. As a relatively new field, the structural biology of MCPs is an area of growing scientific and bioengineering interest [45–56]. Not only a tool for experts in the field, the MCPdb provides novices and even younger students the opportunity to learn about and explore bacterial MCPs. The exceptional biological role of MCPs as protein-based organelles makes them an attractive subject for young scientists, as they challenge the textbook paradigm that eukaryotic cells possess mechanistically complex subcellular organelles while bacterial cells do not.

As the body of structural data on MCPs grows, increased automation will be required to keep the database current. Ongoing developments will involve methods to periodically survey the PDB for new microcompartment and encapsulin-related structures, and their associated data files. Additionally, further efforts will expand the types of information and utilities available on the database. Subsequent versions will introduce an interactive operon map for exploring the operon structure and genomic context of BMC shell protein genes and their associated encapsulated enzymes. We are also working to provide users with further geometric representations of the structures, electrostatic potentials, pore properties, and graphics files for 3D printing. These capabilities will further facilitate access to the field of MCPs for basic and applied research.

## Acknowledgements

The authors thank Alex Lisker, Duilio Cascio, Michael Sawaya, and Kyle Meador for help and advice. This work was supported by NIH/NIAID Grant R01AI081146-08. Jessica M. Ochoa is a Howard Hughes Medical Institute Gilliam Fellow.

## REFERENCES

1. Jorda J, Lopez D, Wheatley NM, Yeates TO. Using comparative genomics to uncover new kinds of protein-based metabolic organelles in bacteria. Protein Science. 2013;22: 179–195. doi:10.1002/pro.2196

2. AbdulRahman F. The Distribution of Polyhedral Bacterial Microcompartments Suggests Frequent Horizontal Transfer and Operon Reassembly. Journal of Phylogenetics & Evolutionary Biology. 2013;01. doi:10.4172/2329-9002.1000118

3. Chowdhury C, Sinha S, Chun S, Yeates TO, Bobik TA. Diverse Bacterial Microcompartment Organelles. Microbiol Mol Biol Rev. 2014;78: 438–468. doi:10.1128/MMBR.00009-14

4. Kerfeld CA, Heinhorst S, Cannon GC. Bacterial microcompartments. Annu Rev Microbiol. 2010;64: 391–408. doi:10.1146/annurev.micro.112408.134211

5. Axen SD, Erbilgin O, Kerfeld CA. A Taxonomy of Bacterial Microcompartment Loci Constructed by a Novel Scoring Method. PLOS Computational Biology. 2014;10: e1003898. doi:10.1371/journal.pcbi.1003898

6. Ravcheev DA, Moussu L, Smajic S, Thiele I. Comparative Genomic Analysis Reveals Novel Microcompartment-Associated Metabolic Pathways in the Human Gut Microbiome. Front Genet. 2019;10. doi:10.3389/fgene.2019.00636

7. Bobik TA, Lehman BP, Yeates TO. Bacterial microcompartments: widespread prokaryotic organelles for isolation and optimization of metabolic pathways. Molecular Microbiology. 2015;98: 193–207. doi:10.1111/mmi.13117

8. Kerfeld CA, Erbilgin O. Bacterial microcompartments and the modular construction of microbial metabolism. Trends in Microbiology. 2015;23: 22–34. doi:10.1016/j.tim.2014.10.003

9. Cannon GC, Bradburne CE, Aldrich HC, Baker SH, Heinhorst S, Shively JM. Microcompartments in Prokaryotes: Carboxysomes and Related Polyhedra. Appl Environ Microbiol. 2001;67: 5351–5361. doi:10.1128/AEM.67.12.5351-5361.2001

10. Yeates TO, Kerfeld CA, Heinhorst S, Cannon GC, Shively JM. Protein-based organelles in bacteria: carboxysomes and related microcompartments. Nature Reviews Microbiology. 2008;6: 681–691. doi:10.1038/nrmicro1913

11. Petit E, LaTouf WG, Coppi MV, Warnick TA, Currie D, Romashko I, et al. Involvement of a Bacterial Microcompartment in the Metabolism of Fucose and Rhamnose by Clostridium phytofermentans. PLOS ONE. 2013;8: e54337. doi:10.1371/journal.pone.0054337

12. Thompson MC, Wheatley NM, Jorda J, Sawaya MR, Gidaniyan SD, Ahmed H, et al. Identification of a Unique Fe-S Cluster Binding Site in a Glycyl-Radical Type Microcompartment Shell Protein. J Mol Biol. 2014;426: 3287–3304. doi:10.1016/j.jmb.2014.07.018

13. Lundin AP, Stewart KL, Stewart AM, Herring TI, Chowdhury C, Bobik TA. Genetic Characterization of a Glycyl Radical Microcompartment Used for 1,2-Propanediol Fermentation by Uropathogenic Escherichia coli CFT073. Metcalf WW, editor. Journal of Bacteriology. 2020;202. doi:10.1128/JB.00017-20

14. Zarzycki J, Sutter M, Cortina NS, Erb TJ, Kerfeld CA. In Vitro Characterization and Concerted Function of Three Core Enzymes of a Glycyl Radical Enzyme - Associated Bacterial Microcompartment. Sci Rep. 2017;7. doi:10.1038/srep42757

15. Wheatley NM, Gidaniyan SD, Liu Y, Cascio D, Yeates TO. Bacterial microcompartment shells of diverse functional types possess pentameric vertex proteins. Protein Sci. 2013;22: 660–665. doi:10.1002/pro.2246

16. Mallette E, Kimber MS. Structural and kinetic characterization of (S)-1-amino-2-propanol kinase from the aminoacetone utilization microcompartment of Mycobacterium smegmatis. J Biol Chem. 2018;293: 19909–19918. doi:10.1074/jbc.RA118.005485

17. Mallette E, Kimber MS. Structure and Kinetics of the S -(+)-1-Amino-2-propanol Dehydrogenase from the RMM Microcompartment of Mycobacterium smegmatis. Biochemistry. 2018;57: 3780–3789. doi:10.1021/acs.biochem.8b00464

18. Lurz R, Mayer F, Gottschalk G. Electron microscopic study on the quaternary structure of the isolated particulate alcohol-acetaldehyde dehydrogenase complex and on its identity with the polygonal bodies of Clostridium kluyveri. Arch Microbiol. 1979;120: 255–262. doi:10.1007/BF00423073

19. Heldt D, Frank S, Seyedarabi A, Ladikis D, Parsons JB, Warren MJ, et al. Structure of a trimeric bacterial microcompartment shell protein, EtuB, associated with ethanol utilization in Clostridium kluyveri. Biochem J. 2009;423: 199–207. doi:10.1042/BJ20090780

20. Tanaka S, Kerfeld CA, Sawaya MR, Cai F, Heinhorst S, Cannon GC, et al. Atomic-Level Models of the Bacterial Carboxysome Shell. Science. 2008;319: 1083–1086. doi:10.1126/science.1151458

21. Yeates TO, Thompson MC, Bobik TA. The protein shells of bacterial microcompartment organelles. Current Opinion in Structural Biology. 2011;21: 223–231. doi:10.1016/j.sbi.2011.01.006

22. Yeates TO, Jorda J, Bobik TA. The Shells of BMC-Type Microcompartment Organelles in Bacteria. Journal of Molecular Microbiology and Biotechnology. 2013;23: 290–299. doi:10.1159/000351347

23. Kerfeld CA, Sawaya MR, Tanaka S, Nguyen CV, Phillips M, Beeby M, et al. Protein Structures Forming the Shell of Primitive Bacterial Organelles. Science. 2005;309: 936–938. doi:10.1126/science.1113397

24. Tsai Y, Sawaya MR, Cannon GC, Cai F, Williams EB, Heinhorst S, et al. Structural Analysis of CsoS1A and the Protein Shell of the Halothiobacillus neapolitanus Carboxysome. PLoS Biol. 2007;5. doi:10.1371/journal.pbio.0050144

25. Yeates TO, Crowley CS, Tanaka S. Bacterial Microcompartment Organelles: Protein Shell Structure and Evolution. Annual Review of Biophysics. 2010;39: 185–205. doi:10.1146/annurev.biophys.093008.131418

26. Bobik TA, Xu Y, Jeter RM, Otto KE, Roth JR. Propanediol utilization genes (pdu) of Salmonella typhimurium: three genes for the propanediol dehydratase. J Bacteriol. 1997;179: 6633–6639. doi:10.1128/jb.179.21.6633-6639.1997

27. Beeby Morgan, Bobik Thomas A., Yeates Todd O. Exploiting genomic patterns to discover new supramolecular protein assemblies. Protein Science. 2008;18: 69–79. doi:10.1002/pro.1

28. Tanaka S, Sawaya MR, Yeates TO. Structure and Mechanisms of a Protein-Based Organelle in Escherichia coli. Science. 2010;327: 81–84. doi:10.1126/science.1179513

29. Crowley CS, Sawaya MR, Bobik TA, Yeates TO. Structure of the PduU Shell Protein from the Pdu Microcompartment of Salmonella. Structure. 2008;16: 1324–1332. doi:10.1016/j.str.2008.05.013

30. Crowley CS, Cascio D, Sawaya MR, Kopstein JS, Bobik TA, Yeates TO. Structural Insight into the Mechanisms of Transport across the Salmonella enterica Pdu Microcompartment Shell. J Biol Chem. 2010;285: 37838–37846. doi:10.1074/jbc.M110.160580

31. Ochoa JM, Nguyen VN, Nie M, Sawaya MR, Bobik TA, Yeates TO. Symmetry breaking and structural polymorphism in a bacterial microcompartment shell protein for choline utilization. Protein Science. 2020;29: 2201–2212. doi:10.1002/pro.3941

32. Sutter M, Greber B, Aussignargues C, Kerfeld CA. Assembly principles and structure of a 6.5-MDa bacterial microcompartment shell. Science. 2017;356: 1293–1297. doi:10.1126/science.aan3289

33. Cai F, Sutter M, Cameron JC, Stanley DN, Kinney JN, Kerfeld CA. The Structure of CcmP, a Tandem Bacterial Microcompartment Domain Protein from the β-Carboxysome, Forms a Subcompartment Within a Microcompartment. J Biol Chem. 2013;288: 16055–16063. doi:10.1074/jbc.M113.456897

34. Sagermann M, Ohtaki A, Nikolakakis K. Crystal structure of the EutL shell protein of the ethanolamine ammonia lyase microcompartment. Proc Natl Acad Sci U S A.2009;106: 8883–8887. doi:10.1073/pnas.0902324106

35. Thompson MC, Cascio D, Leibly DJ, Yeates TO. An allosteric model for control of pore opening by substrate binding in the EutL microcompartment shell protein. Protein Science. 2015;24: 956–975. doi:10.1002/pro.2672

36. Takenoya M, Nikolakakis K, Sagermann M. Crystallographic Insights into the Pore Structures and Mechanisms of the EutL and EutM Shell Proteins of the Ethanolamine-Utilizing Microcompartment of Escherichia coli. Journal of Bacteriology. 2010;192: 6056–6063. doi:10.1128/JB.00652-10

37. Pang A, Warren MJ, Pickersgill RW. Structure of PduT, a trimeric bacterial microcompartment protein with a 4Fe–4S cluster-binding site. Acta Cryst D, Acta Cryst Sect D, Acta Crystallogr D, Acta Crystallogr Sect D, Acta Crystallogr D Biol Crystallogr, Acta Crystallogr Sect D Biol Crystallogr. 2011;67: 91–96. doi:10.1107/S0907444910050201

38. Mallette E, Kimber MS. Structural and kinetic characterization of (S)-1-amino-2-propanol kinase from the aminoacetone utilization microcompartment of Mycobacterium smegmatis. J Biol Chem. 2018; jbc.RA118.005485. doi:10.1074/jbc.RA118.005485

39. Keeling TJ, Samborska B, Demers RW, Kimber MS. Interactions and structural variability of β-carboxysomal shell protein CcmL. Photosynth Res. 2014;121: 125–133. doi:10.1007/s11120-014-9973-z

40. Burley SK, Berman HM, Kleywegt GJ, Markley JL, Nakamura H, Velankar S. Protein Data Bank (PDB): The Single Global Macromolecular Structure Archive. In: Wlodawer A, Dauter Z, Jaskolski M, editors. Protein Crystallography: Methods and Protocols. New York, NY: Springer; 2017. pp. 627–641. doi:10.1007/978-1-4939-7000-1_26

41. Consortium TU. UniProt: a worldwide hub of protein knowledge. Nucleic Acids Res. 2019;47: pD506–D515. doi:10.1093/nar/gky1049

42. Krissinel E, Henrick K. Inference of Macromolecular Assemblies from Crystalline State. Journal of Molecular Biology. 2007;372: 774–797. doi:10.1016/j.jmb.2007.05.022

43. Niknafs N, Kim D, Kim R, Diekhans M, Ryan M, Stenson PD, et al. MuPIT interactive: webserver for mapping variant positions to annotated, interactive 3D structures. Hum Genet. 2013;132: 1235–1243. doi:10.1007/s00439-013-1325-0

44. Carrillo-Tripp M, Shepherd CM, Borelli IA, Venkataraman S, Lander G, Natarajan P, et al. VIPERdb2: an enhanced and web API enabled relational database for structural virology. Nucleic Acids Research. 2009;37: pD436–D442. doi:10.1093/nar/gkn840

45. Parsons JB, Frank S, Bhella D, Liang M, Prentice MB, Mulvihill DP, et al. Synthesis of Empty Bacterial Microcompartments, Directed Organelle Protein Incorporation, and Evidence of Filament-Associated Organelle Movement. Molecular Cell. 2010;38: 305–315. doi:10.1016/j.molcel.2010.04.008

46. Lawrence AD, Frank S, Newnham S, Lee MJ, Brown IR, Xue W-F, et al. Solution structure of a bacterial microcompartment targeting peptide and its application in the construction of an ethanol bioreactor. ACS Synth Biol. 2014;3: 454–465. doi:10.1021/sb4001118

47. Huber I, Palmer DJ, Ludwig KN, Brown IR, Warren MJ, Frunzke J. Construction of Recombinant Pdu Metabolosome Shells for Small Molecule Production in Corynebacterium glutamicum. ACS Synth Biol. 2017;6: 2145–2156. doi:10.1021/acssynbio.7b00167

48. Wagner HJ, Capitain CC, Richter K, Nessling M, Mampel J. Engineering bacterial microcompartments with heterologous enzyme cargos. Engineering in Life Sciences. 2017;17: 36–46. doi:https://doi.org/10.1002/elsc.201600107

49. Fang Y, Huang F, Faulkner M, Jiang Q, Dykes GF, Yang M, et al. Engineering and Modulating Functional Cyanobacterial CO2-Fixing Organelles. Front Plant Sci. 2018;9. doi:10.3389/fpls.2018.00739

50. Hagen A, Sutter M, Sloan N, Kerfeld CA. Programmed loading and rapid purification of engineered bacterial microcompartment shells. Nature Communications. 2018;9: 2881. doi:10.1038/s41467-018-05162-z

51. Lee MJ, Mantell J, Brown IR, Fletcher JM, Verkade P, Pickersgill RW, et al. De novo targeting to the cytoplasmic and luminal side of bacterial microcompartments. Nature Communications. 2018;9: 3413. doi:10.1038/s41467-018-05922-x

52. Huang J, Ferlez BH, Young EJ, Kerfeld CA, Kramer DM, Ducat DC. Functionalization of Bacterial Microcompartment Shell Proteins With Covalently Attached Heme. Frontiers in Bioengineering and Biotechnology. 2020;7. doi:10.3389/fbioe.2019.00432

53. Jakobson CM, Chen Y, Slininger MF, Valdivia E, Kim EY, Tullman-Ercek D. Tuning the Catalytic Activity of Subcellular Nanoreactors. Journal of Molecular Biology. 2016;428: 2989–2996. doi:10.1016/j.jmb.2016.07.006

54. Quin MB, Perdue SA, Hsu S-Y, Schmidt-Dannert C. Encapsulation of multiple cargo proteins within recombinant Eut nanocompartments. Appl Microbiol Biotechnol. 2016;100: 9187–9200. doi:10.1007/s00253-016-7737-8

55. Lee MJ, Mantell J, Hodgson L, Alibhai D, Fletcher JM, Brown IR, et al. Engineered synthetic scaffolds for organizing proteins within the bacterial cytoplasm. Nature Chemical Biology. 2018;14: 142–147. doi:10.1038/nchembio.2535

56. Zhang G, Johnston T, Quin MB, Schmidt-Dannert C. Developing a Protein Scaffolding System for Rapid Enzyme Immobilization and Optimization of Enzyme Functions for Biocatalysis. ACS Synthetic Biology. 2019;8: 1867–1876. doi:10.1021/acssynbio.9b00187

